# Reconstructing living materials as a computable design space with multi-agent reasoning

**DOI:** 10.64898/2026.02.15.705954

**Authors:** Yewei Xiao, Xi Zeng, Zhaoxiang Yang, Junwei Gu, Yanhong Lu, Yujie Wang, Huilin Wen, Mingdong Chen, Zhipeng Huang, Junwei Hu, Jiaojiao Liu, Chengxiang Sha, Jinhui Xie, Huan Li, Xiang Zhu, Shenchi Zheng, Jing Zhang, Weijie Zong, Zehao He, Ying Xu, Xin Zhou, Fei Li, Hui Liu, Qiaoning He, Linlin Liu, Ziyi Yu

**Author notes:** These authors are equal contributed to this work (Yewei Xiao, Xi Zeng, Zhaoxiang Yang).

## Abstract

Artificial intelligence is increasingly used to accelerate scientific discovery, but most successful frameworks operate within well-defined molecular, protein or materials spaces. Living materials present a more formidable computational problem because functions emerge from context dependent coupling among cells, matrices, fabrication processes and evaluation conditions. Here we introduce LiveMat, a multi-agent reasoning framework that transforms unstructured literature into a computable design space for living materials. LiveMat standardizes 34,215 living material records, integrating 16,769 microorganism and 17,446 polymer entries into a knowledge graph linking living components, abiotic matrices, functional outputs, evaluation contexts and performance metrics. Benchmarking across five large language models shows that living material reasoning is limited mainly by cross-domain feature integration rather than coarse classification. LiveMat overcomes this limitation through constraint decomposition, provenance-aware extraction, consistency checking and expert-anchored ranking. In a prospective wound-healing task, it prioritizes a four-component design with state-of-the-art *in vivo* performance, establishing a scalable infrastructure for interpretable, evidence-grounded living material discovery.

## Introduction

Advancements in multidisciplinary scientific design hinge not on the availability of candidate components, but on the creation of computable representations that can convert fragmented knowledge into actionable design rules^1-10^. Living materials are an important class of biohybrid systems^11-14^ because they integrate living cells or microorganisms with abiotic matrices to generate adaptive^15^, regenerative^16-17^, sensing^18-20^, manufacturing^21-27^ and environmental functions^28-30^ that are inaccessible to inert materials alone. Their potential spans therapeutic delivery^16^, tissue repair^17^, sustainable biomanufacturing^21-24^, environmental remediation^28-30^ and responsive devices^18-20^, yet their design remains difficult because function is not encoded in cells or matrices independently. Instead, performance emerges from context-dependent coupling among living components, abiotic matrices, fabrication histories and local microenvironments^32-35^. Rational design therefore requires a framework that can preserve these cross-domain constraints and reason over biological feasibility, material function, processability and application specific performance.

The knowledge required for such reasoning is abundant but weakly formalized. Experimental reports often describe microbial chassis, matrix chemistry, fabrication conditions, biological outputs and performance metrics in narrative form, with limited standardization across studies. Negative results, failed combinations and boundary conditions are rarely encoded as reusable design constraints^37-39^. Feasibility, compatibility, biosafety and processability are therefore often implicit assumptions rather than explicit variables. Existing reviews and databases can summarize components or applications, but they rarely capture the relational design logic^37-39^ that links cellular traits, material properties, processing variables and functional outcomes^32-35^. Similarly, general-purpose language models can retrieve and restate relevant information^1-5^, but their lack of explicit design representations often leads them to violate critical domain constraints, overgeneralize from partial evidence or generate plausible but unsupported combinations^1-10^.

Recent advances in artificial intelligence (AI) create an opportunity to extract, structure and reason over unstructured scientific literature^1-9^. However, most AI-driven discovery frameworks remain grounded in relatively well-defined molecular, protein or materials spaces, where the object of design can be represented as a molecule, sequence, structure or composition^5-10^. Living materials require a different formulation, in which the design object is a context-dependent living material configuration rather than an isolated molecule, polymer or cell^11-42^. They therefore provide a stringent test case for extending AI for Science from single domain prediction to auditable cross-domain reasoning^43-44^. Here we introduce LiveMat, a retrieval-grounded multi-agent reasoning framework that reconstructs living materials from unstructured literature into a computable design space. LiveMat converts document-level knowledge into a domain-scale design graph linking living components, abiotic matrices, fabrication strategies, functional outputs, evaluation contexts and performance metrics. It combines knowledge-construction agents with design-reasoning agents to support evidence-grounded retrieval, cross-domain constraint integration, expert-anchored evaluation and negative-constraint feedback. We use LiveMat to distill literature-scale design priors, benchmark cross-domain reasoning failures in general-purpose large language models, and prospectively validate a constraint-ranked design in an acute wound-healing task. These results establish a route for converting fragmented empirical knowledge into cumulative, auditable and experimentally grounded design rules for living materials.

## Results

### Reconstructing living materials as a computable design space

To formalize living material design as a machine actionable problem, LiveMat represents each reported system as a relational configuration of biological, material, and evaluation variables (Fig. 1A). This formulation supersedes traditional approaches that treat microorganisms and polymers as isolated descriptors. Instead, each living material is encoded through coupled constraints (Fig. 2A-D), including microbial viability, material compatibility, processing feasibility, and application-specific requirements. By formalizing these interdependencies, LiveMat converts living material design from an experience driven search over individual components into a structured design problem that can be queried, compared, ranked and updated across studies.

**Figure 1.**
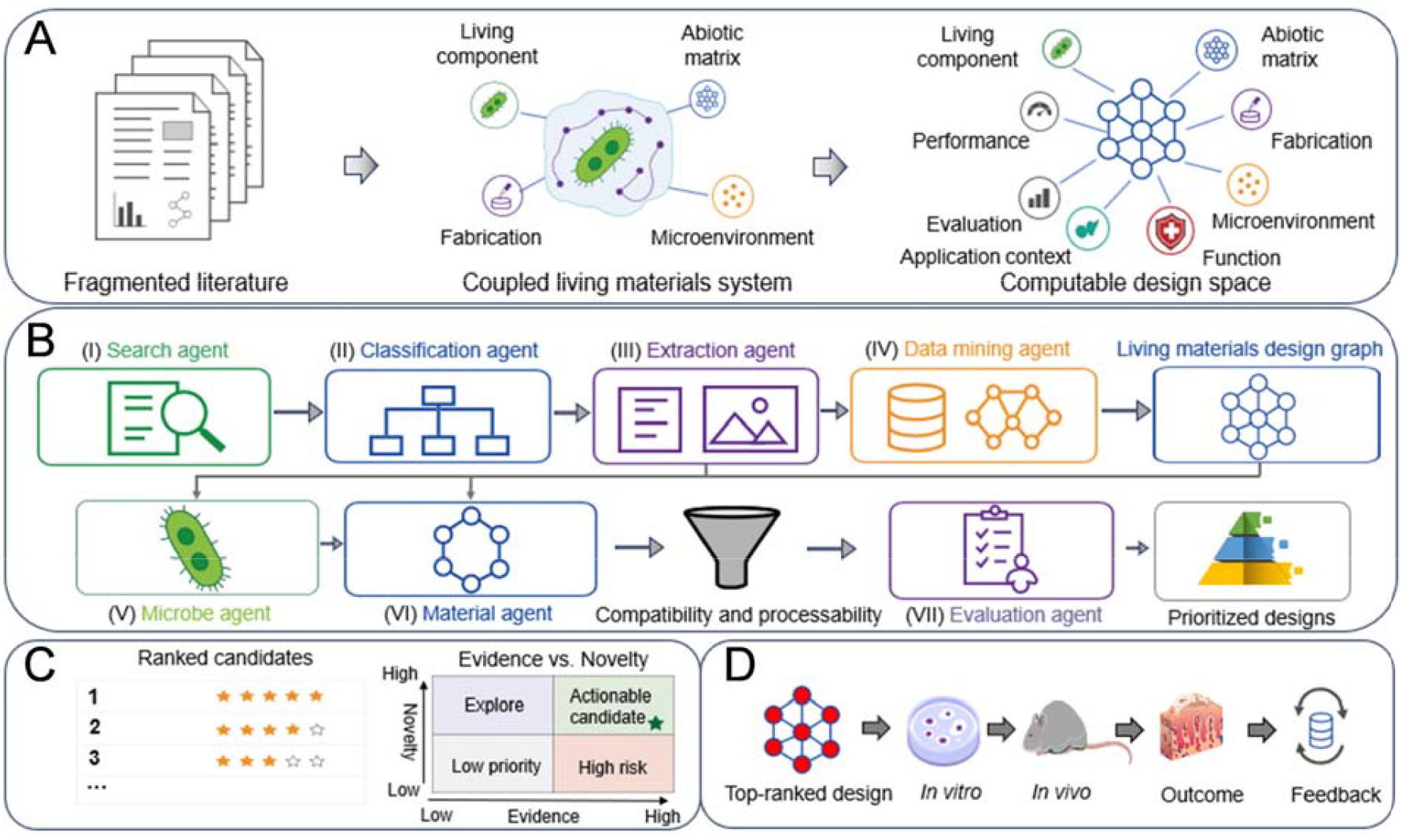
LiveMat reconstructs fragmented living materials knowledge into a computable multi-agent design space. (A) Fragmented literature is converted into a structured representation linking cells, matrices, fabrication processes, contexts, functions, metrics and outcomes. (B) Seven-agent workflow for evidence construction and design reasoning, integrating literature retrieval, classification, extraction, graph construction, component evaluation, compatibility filtering and candidate ranking. (C) Ranked candidates are mapped within an evidence –novelty space to support downstream selection. (D) Prospective validation links computational design, experimental testing and feedback-driven refinement.

**Figure 2.**
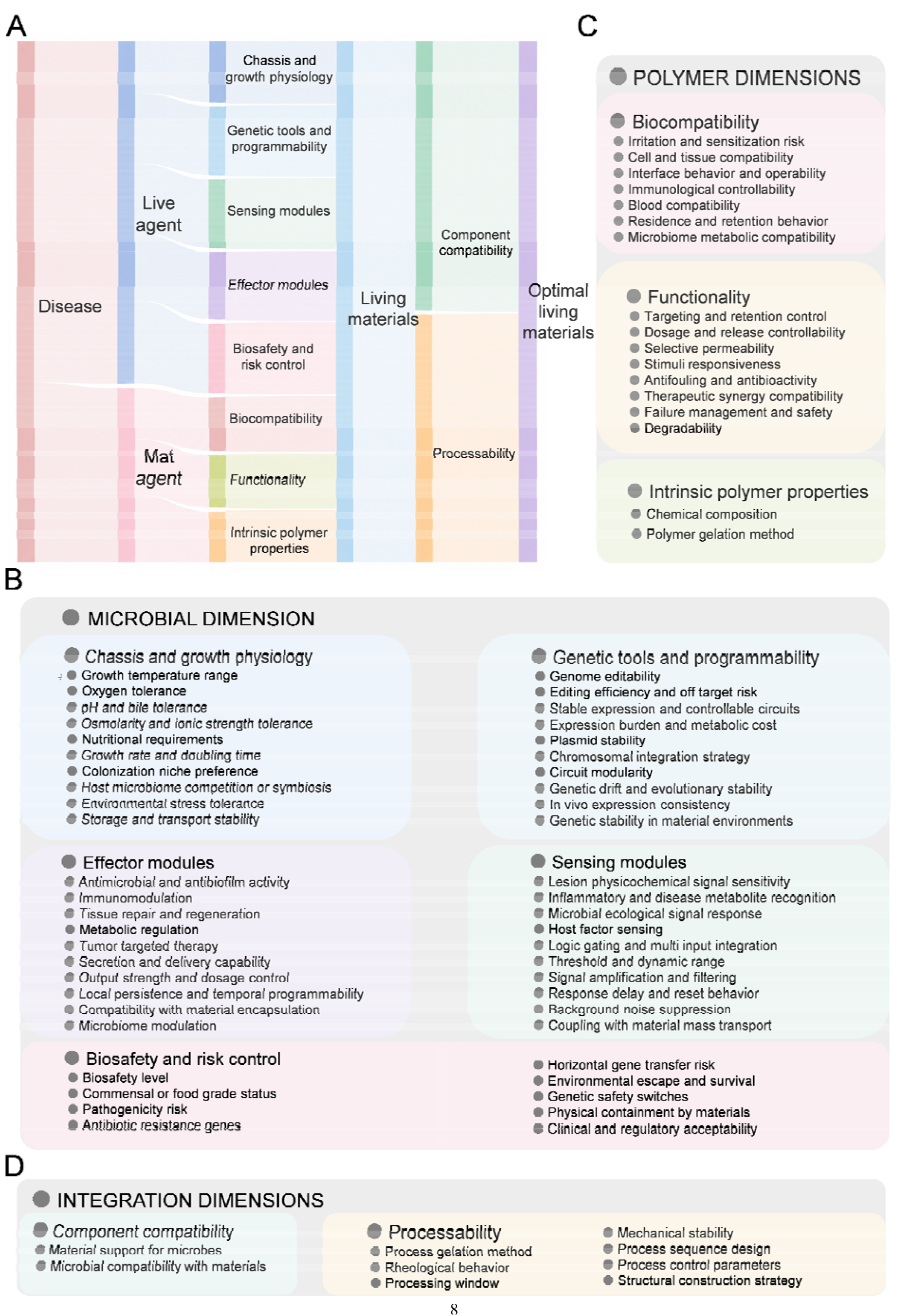
LiveMat ontology decomposes living materials design into microbial, material and integration dimensions. (A) Application requirements are decomposed by the Live Agent and Mat Agent into microbial and material design dimensions, followed by compatibility and processability reasoning. (B) Microbial ontology covering chassis physiology, genetic programmability, effector functions, sensing modules and biosafety control. (C) Material ontology covering biocompatibility, functionality and intrinsic material properties relevant to biological performance and fabrication. (D) Integration ontology evaluating component compatibility and processability to determine whether microbial and material candidates can form feasible living-material systems.

As illustrated in Fig. 1, LiveMat implements this formulation through a seven-agent workflow (Fig. 1B; Supplementary Fig. S1) that separates knowledge construction from design reasoning. Four knowledge construction agents transform heterogeneous scientific evidence into reusable design information: the Search Agent retrieves relevant literature and database records; the Classification Agent filters records from coarse relevance to fine-grained domain categories; the Extraction Agent parses text- and figure-derived information into structured entities, relations and experimental attributes; and the Data Mining Agent integrates extracted features with curated external databases and graph-level descriptors. These processes generate a living material design graph linking living components, abiotic matrices, fabrication strategies, functional outputs, application contexts and evaluation metrics (Fig. 1C). On this graph, three design reasoning agents support candidate prioritization: The Microbe Agent ranks organisms under biological, functional and biosafety constraints; the Material Agent prioritizes polymers according to physicochemical feasibility, biocompatibility and functional roles; and the Evaluation Agent assesses candidate configurations using multidimensional criteria (Fig. 1D), evidence attribution and expert verification.

The resulting design graph organizes living materials into coordinated microbial and material subspaces (Fig. 2A, B) connected through compatibility and processability filters. The microbial subspace captures chassis and growth physiology, genetic tools and programmability, sensing modules, effector modules, and biosafety and risk control, whereas the material subspace captures intrinsic polymer properties, material functionality and biocompatibility. Candidate living materials are generated by coupling these subspaces (Fig. 2C, D) through a constraint layer that evaluates microenvironmental support for microbial survival and function, cellular tolerance to matrix and fabrication conditions, and the experimental and translational feasibility of the resulting configurations. Evidence provenance, intermediate reasoning steps, expert annotations and evaluation outcomes are retained within the design graph, while unsupported claims and unsuccessful candidates are recorded as negative constraints for subsequent reasoning cycles. This computable representation provides the foundation for literature scale reconstruction of design priors, benchmarking of cross-domain reasoning and constraint-driven candidate ranking in the following sections.

### Feature-level benchmarking enables literature-scale reconstruction of living-material design priors

To assess whether the LiveMat could support reliable reasoning across heterogeneous living material tasks, we first benchmarked five large language models (Fig. 3A) under a shared evaluation framework. The benchmark covered response time, token consumption (Supplementary Fig. S3), classification accuracy and feature-level reasoning across microorganism, polymer and integrated living-material dimensions. Although classification level accuracy was high across models, this metric alone provided limited discrimination. Most models could recognize whether a record was relevant to microorganisms, polymers or living materials, indicating that coarse domain classification is not the main bottleneck in living-material reasoning.

**Figure 3.**
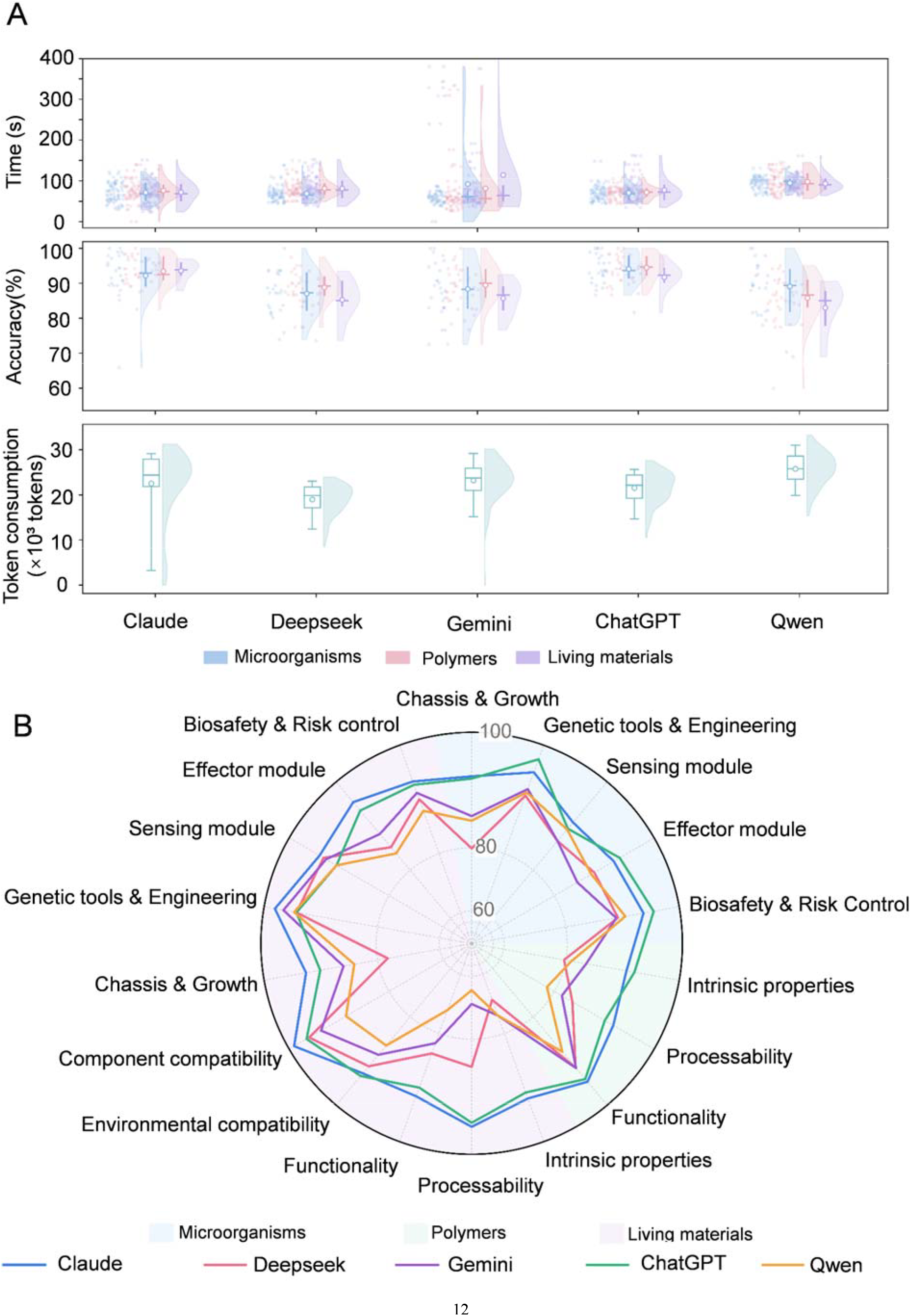
Benchmarking reveals model efficiency and feature-level accuracy differences among large language models. (A) Benchmarking of five large language models across microorganism, material and living-material design tasks. Raincloud plots compare response time and accuracy, whereas aggregate token consumption indicates model-level computational cost. (B) Radar plots showing feature-level accuracy across microbial, material and living-material dimensions. Model differences are most pronounced in structured feature integration, highlighting the need for multidimensional evaluation beyond coarse task accuracy.

Feature-level evaluation revealed a different pattern. Model performance diverged substantially (Fig. 3B; Supplementary Figs. S4 and S5) when the task required extraction and integration of fine-grained biological and material constraints. GPT-5.4 showed the most balanced performance across microorganism, polymer and living material axes, whereas other models exhibited greater variation, particularly in recall and in dimensions requiring cross-domain interpretation. Precision–recall analysis further indicated that many errors arose from omitted constraints rather than incorrect statements. These results show that the major challenge for AI-assisted living material design lies not in recognizing domain relevance, but in preserving and integrating feature level constraints across biological, material and application contexts.

We then applied the benchmarked LiveMat workflow to reconstruct a literature-scale living material design landscape. Across the reconstructed corpus, LiveMat standardized 34,215 living material records, comprising 16,769 microorganism entries and 17,446 polymer entries (Fig. 4B). Figure 4A separately summarizes the literature-frequency distribution of these records across microorganism and polymer categories, yielding 34,738 category-frequency counts in total, including 16,056 microorganism-associated counts and 18,682 polymer-associated counts. This reconstruction revealed strong design priors in the field. Bacterial chassis dominated the biological space, whereas natural polymers dominated the material space. Frequently used organisms such as *Lactobacillus, Bacillus* and *Escherichia*, together with materials such as alginate, chitosan, gelatin and polyethylene glycol, formed high-density regions within the design landscape (Supplementary Figs. S6-S9).

**Figure 4.**
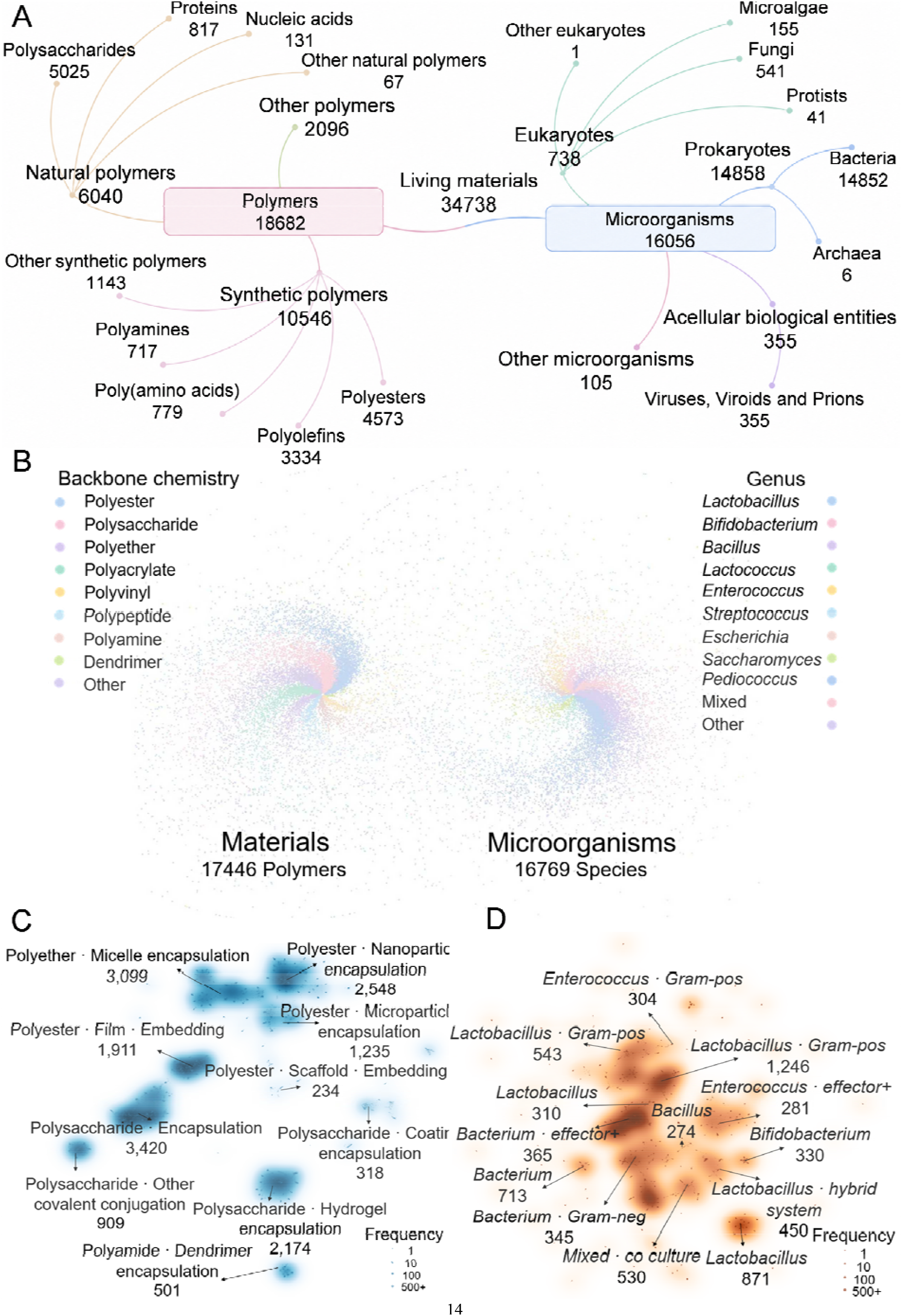
Literature-scale reconstruction reveals the composition, organization and density structure of the living materials design space. (A) Hierarchical organization of the reconstructed corpus into material and microorganism subspaces. Numbers indicate literature/category-frequency counts for each branch rather than standardized entry counts. (B) UMAP map of standardized polymer and microorganism entries reconstructed by LiveMat, colored by material chemistry or microbial taxonomy. (C) Density landscape of the material subspace reconstructed from standardized polymer entries, highlighting dominant formulation paradigms and underexplored material regions. (D) Density landscape of the microorganism subspace reconstructed from standardized microorganism entries, revealing frequently studied microbial taxa and sparse regions for future exploration.

The resulting knowledge graph further showed that living material research is concentrated around recurrent microorganism-material pairings (Fig. 4C, D). These dense regions represent experimentally familiar and translationally conservative design choices, whereas sparsely connected regions point to underexplored combinations with potential discovery value (Supplementary Figs. S10-S13 and S14A-D). By converting fragmented literature into a structured and evaluable design landscape, LiveMat makes the field’s implicit design priors explicit and provides a foundation for both conservative and exploratory living-material design.

### Constraint-driven reasoning identifies a top ranked living material design

To test whether LiveMat can perform prospective design beyond retrospective literature reconstruction, we formulated wound-healing living material design as a constrained four-component reasoning task. Rather than merely generating a plausible formulation, the objective was to evaluate whether LiveMat could systematically translate application-level therapeutic requirements into explicit biological and material constraints, and subsequently rank traceable living-material configurations (Fig. 5A). From the unstructured requirement, LiveMat identified four high priority constraints after expert confirmation: antibacterial activity and oxygen production on the microorganism side, corresponding to infection suppression and hypoxia mitigation; and interface stabilization and therapeutic support on the material side, corresponding to maintenance of a hydrated wound interface and enrichment of regenerative cues. These constraints were converted into explicit screening rules (Fig. 5A) for downstream candidate search, ranking and cross-domain evaluation.

**Figure 5.**
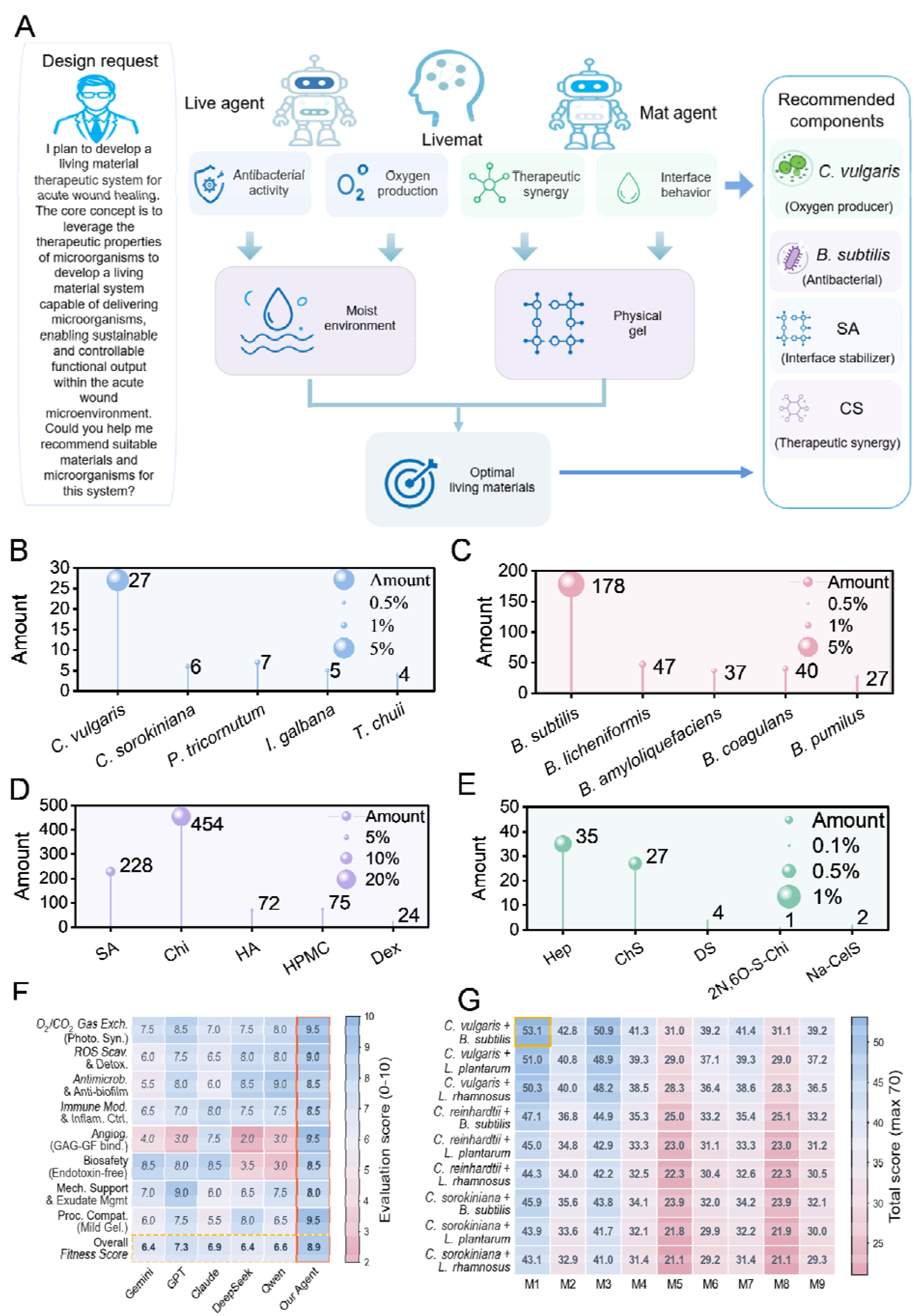
Multi-agent constraint reasoning identifies and prioritizes a four-component living materials design for acute wound healing. (A) LiveMat decomposes a wound-healing request into microbial, material and integration constraints, leading to a four-component design comprising Chlorella vulgaris, Bacillus subtilis, sodium alginate and chondroitin sulfate. (B, C) Ranked oxygen-producing and antibacterial microorganism candidates identified by the Live Agent. (D, E) Ranked material candidates associated with therapeutic synergy, interface regulation and matrix stabilization. (F) Multi-criterion comparison of LiveMat- and LLM-generated designs across wound-healing-relevant biological, material and processing criteria. (G) Combinatorial fitness landscape showing that the LiveMat-prioritized system occupies the highest-scoring design region under explicit biological and material constraints.

Candidate selection was then performed within agent-specific design spaces (Fig. 5B-E) rather than by direct formulation generation. In the Live Agent space, microorganism candidates were evaluated across chassis and growth physiology, genetic tools and engineering plasticity, sensing modules, effector modules, and biosafety and containment. Under these criteria, *Bacillus subtilis* (*B. subtilis*) was prioritized as the antibacterial chassis by integrating antimicrobial activity, biosafety profile, metabolic compatibility and evidence density, whereas *Chlorella vulgaris* (*C. vulgaris*) was selected from photoautotrophic candidates because it showed consistent support for sustained oxygen production under biologically relevant conditions. In parallel, the Mat Agent evaluated polymer candidates across intrinsic polymer properties, processing and structural properties, and functionality. Sodium alginate (SA) was prioritized for hydrophilicity, gel-forming capability, biocompatibility and microbial encapsulation, whereas chondroitin sulfate (CS) was prioritized for extracellular-matrix-associated signalling and growth-factor interaction. These candidates were then passed into the living material reasoning layer (Fig. 5B-E), where component compatibility and processability windows were evaluated before generating the final four-component design.

This reasoning process yielded a candidate system comprising *B. subtilis, C. vulgaris*, SA and CS. Importantly, this configuration was not obtained by manual enumeration or by asking a language model to directly generate a formulation. It emerged from the alignment of agent-led requirement decomposition, subspace-specific candidate ranking, living material compatibility assessment and processability evaluation, making each component choice traceable to an explicit reasoning step. To assess whether the same design could be recovered without this structured reasoning process, we compared LiveMat with direct LLM prompting as a task-level baseline. Five general-purpose large language models were given the same four-component design task and evaluated using a standardized scoring framework across eight functional criteria (Fig. 5F), including antimicrobial activity, oxygen production, angiogenesis support, biosafety, mechanical support and process compatibility.

Direct prompting produced superficially plausible living material systems, but the resulting designs showed uneven constraint satisfaction. Some models emphasized antimicrobial activity while introducing microorganisms with weaker biosafety or endotoxin-related concerns, whereas others selected hydrogel matrices with acceptable general biocompatibility but weaker support for microbial co-encapsulation, growth factor interaction or fabrication compatibility. By contrast, the LiveMat-ranked design achieved the highest overall score (Supplementary Fig. S15) while maintaining consistently high scores across biological, material and therapeutic criteria (Fig. 5F), balancing antibacterial activity, oxygen production, hydrated interface maintenance, therapeutic support, biocompatibility, component compatibility and processability. To systematically determine whether this result reflected a robust ranking rather than a single generated answer, we constructed a 9 × 9 combinatorial design matrix by pairing top-ranked antibacterial and oxygen-producing microbial modules with top-ranked material modules. The *B. subtilis*-*C. vulgaris*-SA–CS configuration occupied the highest scoring region of this explicitly enumerated landscape (Fig. 5G), indicating that LiveMat identified not merely a plausible formulation, but a balanced and traceable design within a constrained living-material design space. These results show that LiveMat does not simply generate plausible living material combinations. Instead, it performs constraint-driven ranking across biological and material subspaces, identifies trade-offs among candidate systems and selects a balanced four component design that outperforms direct LLM-generated alternatives.

### Experimental validation confirms the predicted living material design

Following the identification of the top-ranked four-component configuration by constraint-driven reasoning, we next designed an experimental validation workflow to examine whether the LiveMat predicted system could be physically realized and whether the resulting living material could improve wound repair *in vivo* (Fig. 5A). To preserve the two predicted microbial functions within a processable SA/CS matrix, *B. subtilis*- or *C. vulgaris*-laden methacrylated alginate microgels were first generated by calcium crosslinking and mechanical fragmentation, and then incorporated into a SA/CS precursor to form a 405 nm-photocrosslinkable composite bioink (Fig. 6A) (Fig. S16A). This fabrication strategy produced a dual-crosslinked hydrogel system in which Ca^2^ □ -mediated ionic crosslinking supported injectability and shear-thinning processability, whereas photoinduced covalent crosslinking stabilized the printed architecture (Fig. S16B). As shown in Fig. S16C-E, rheological analyses confirmed rapid network reinforcement after light irradiation and enhanced elastic stability in the dual-crosslinked hydrogels compared with single-crosslinked controls, consistent with the processability and mechanical constraints encoded in the LiveMat design. Microscopic imaging further showed that *B. subtilis* and *C. vulgaris* were not homogeneously mixed at the single-cell level, but remained spatially compartmentalized within microorganism-specific microgel domains (Fig. S17A). This organization preserved local microbial niches while integrating the two functional species within one continuous hydrogel network. Scanning electron microscopy further revealed distinct microbial entities embedded in a porous hydrogel matrix, confirming successful co-embedding and structural integration of the LiveMat-predicted biological components (Fig. S17B).

**Figure 6.**
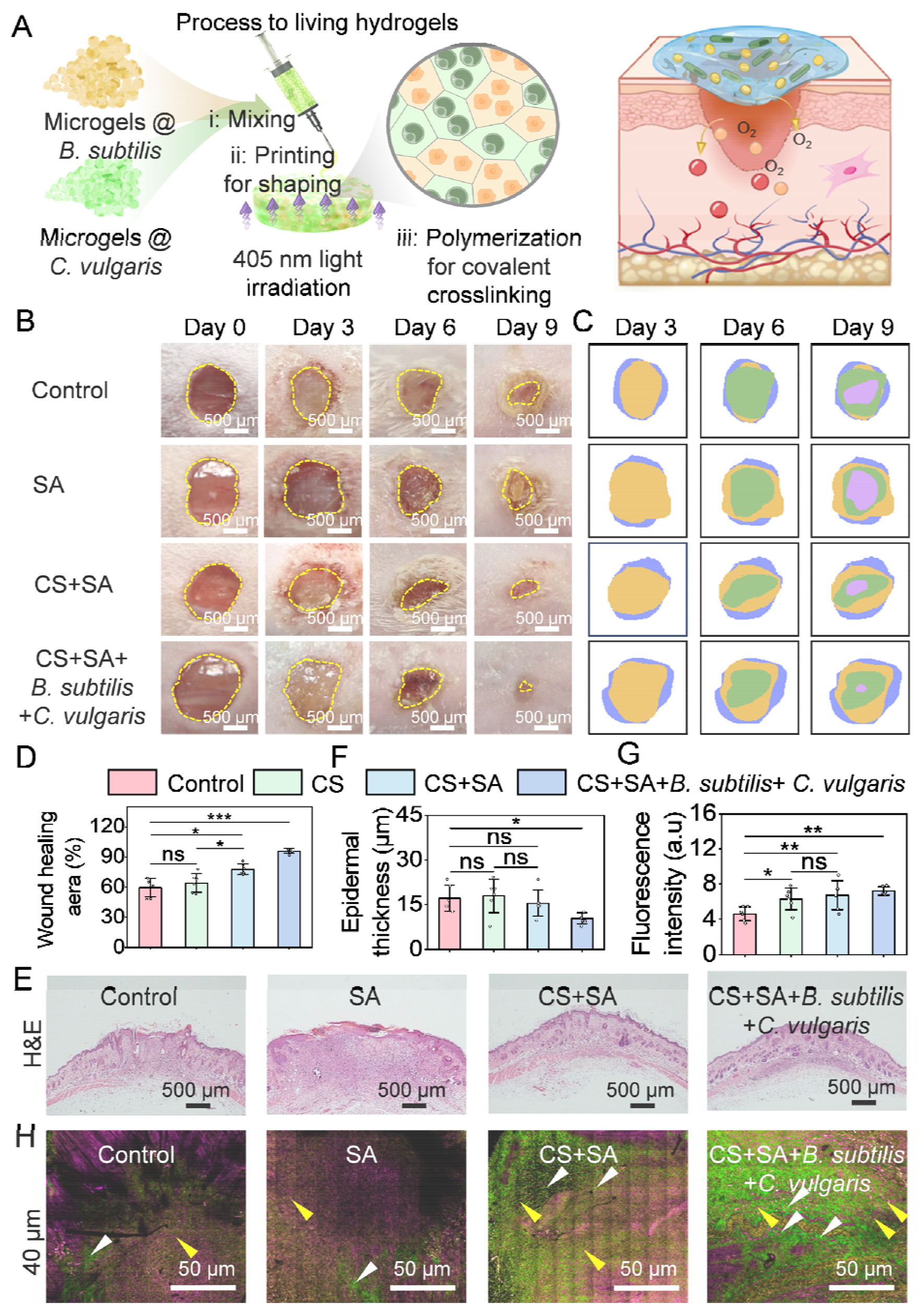
Fabrication and *in vivo* validation of the living-material system. (A) Fabrication scheme of the living hydrogel by microgel mixing, printing-assisted shaping and photocrosslinking. (B-D) Representative images and quantitative analysis of wound closure in different groups on days 0, 3, 6, and 9 after live material system treatment. (E, F) Representative H&E staining images and quantitative analysis of epidermal thickness in wound tissues from different groups. (G) Quantitative analysis of the fluorescence intensity of CD31 in the wound tissues. (H) Representative femtosecond label-free imaging showing collagen (white arrow) and fibroblasts (yellow arrow). Data are presented as mean ± S.D. (n = 6, n represents biological replicates). \**P* < 0.05, \*\**P* < 0.01, and \*\*\**P* < 0.001 vs. the control group.

We then evaluated whether the physically realized living material could promote wound repair *in vivo*. In the full-thickness dorsal wound model, wound areas gradually decreased in all groups, but the healing dynamics differed markedly among treatments. Compared with the control group, wounds treated with the CS + SA + *B. subtilis* + *C. vulgaris* living material showed smaller residual wound areas and faster wound contraction throughout the healing process. Quantitative analysis further confirmed a significantly higher wound-closure rate in the four component group (Fig. 6B-D). Notably, the CS + SA matrix promoted wound healing more effectively than SA alone, supporting the LiveMat-predicted role of CS in therapeutic support, likely through enrichment of regenerative cues and stabilization of the wound microenvironment (Fig. 6B-D).

To clarify the tissue level basis of this improved healing response, we performed histological, immunofluorescence and label-free imaging analyses. H&E staining showed that the newly formed epidermis in the CS + SA + *B. subtilis* + *C. vulgaris* group was more continuous and complete than that in the control group, with increased epidermal thickness (Fig. 6E, F). CD31 immunofluorescence staining revealed stronger angiogenic activity in the four-component group, as indicated by enhanced CD31-positive signals (Fig. S18A, Fig. 6G). Femtosecond label-free imaging further showed pronounced fibroblast infiltration and a dense, interconnected collagen network in treated wounds, indicating enhanced extracellular matrix remodeling (Fig. 6H). Systemic biosafety evaluation showed no detectable pathological abnormalities in major organs and stable body weight across groups (Fig. S18B-G). Collectively, these results confirm that the LiveMat-predicted four-component system can be translated into a compartmentalized living material that accelerates wound closure and promotes coordinated tissue regeneration while maintaining a favorable biosafety profile.

## Discussion and conclusion

We have developed LiveMat as a multi-agent reasoning framework for reconstructing living materials as a computable design space. Unlike direct formulation generation, LiveMat treats living material design as a cross-domain reasoning problem in which organisms, matrices, functional outputs and evaluation contexts must be represented and assessed jointly. This formulation is necessary because the performance of living materials is not determined by a single cell type or component, but emerges from context-dependent interactions among microbial physiology, matrix properties, processing conditions and application-specific microenvironments. By transforming unstructured literature into structured records and a relational design graph, LiveMat provides a machine actionable foundation for querying, comparing and ranking heterogeneous living material configurations.

A central outcome of this work is that living material design can be shifted from component level intuition to constraint-driven reasoning. The reconstruction of 34,738 living material records revealed both the density structure and the implicit priors of the field, including the dominance of bacterial chassis, natural polymers and recurrent microorganism-matrix pairings. These patterns provide useful evidence for conservative design, but they also expose sparsely explored regions where new combinations may be discovered. Benchmarking across five large language models further showed that the main limitation of current AI-assisted living material design is not coarse domain recognition, but the preservation and integration of fine-grained biological, material and process constraints. LiveMat addresses this limitation by decomposing design requirements, preserving evidence provenance, checking cross-domain consistency and ranking candidates through expert-anchored evaluation.

The prospective wound healing task demonstrates the practical utility of this framework. LiveMat translated an open-ended therapeutic request into explicit microbial, material and integration constraints, leading to a four-component system composed of *B. subtilis, C. vulgaris*, SA and CS. This design balanced antibacterial activity, oxygen supply, hydrated interface maintenance, therapeutic support, biosafety, component compatibility and processability. Experimental validation confirmed that the predicted system could be physically realized as a compartmentalized living hydrogel and that it accelerated wound closure, enhanced epidermal regeneration, promoted angiogenesis and supported extracellular matrix remodeling in vivo. These results show that LiveMat does not merely generate plausible formulations, but can identify traceable, experimentally actionable designs within a constrained living-material design landscape.

More broadly, LiveMat establishes a scalable route for evidence grounded and interpretable AI-assisted discovery in living materials. Its current implementation focuses on literature derived evidence and expert defined evaluation criteria, but the framework can be expanded by incorporating negative results, high-throughput experimental data, quantitative performance models and closed-loop optimization. Such extensions may allow living material design graphs to evolve from static knowledge repositories into continuously updated discovery engines. By making biological, material and process constraints explicit and reusable, LiveMat provides a generalizable strategy for designing living materials across therapeutic, environmental, sensing and biomanufacturing applications, and offers a model for extending AI for Science from single-domain prediction toward auditable reasoning over complex biohybrid systems.

## Methods

### System architecture and agent roles

LiveMat was implemented as a multi-agent reasoning framework for living materials discovery^1-7^. The system comprised seven functional agents corresponding to literature search, hierarchical classification, multimodal extraction, data mining and graph integration, microbe ranking, material ranking and evaluation. These agents operated under centralized task orchestration and shared a common evidence context. The search agent retrieved literature and database records, the classification agent filtered domain-relevant documents, the extraction agent parsed text- and figure-derived information, the data agent integrated extracted features with curated databases, the microbe agent ranked organisms under biological, functional and biosafety constraints, the material agent ranked polymers according to physicochemical, biocompatibility and functional criteria, and the evaluation agent assessed candidate outputs against expert-defined criteria.

Each agent performed a constrained function rather than generating final designs independently. Intermediate evidence, extracted features, reasoning traces and evaluation outcomes were stored in a structured database and linked to a domain-scale knowledge graph^2-3^. This design allowed subsequent reasoning steps to use both positive evidence and recorded failure modes.

### Large language model configuration

The multi-LLM reasoning core coordinated multiple large language models under fixed generation settings^1-5^. The five evaluated models were Gemini-3.1-Pro, ChatGPT-5.4, Claude-opus-4.6, DeepSeek-V3.2 and Qwen-3.5-Plus (Fig. 3A; Supplementary Figs. S3-S5). Unless otherwise specified, temperature and sampling parameters were kept constant across tasks to reduce stochastic variation. Prompts were defined before evaluation and were not modified during benchmark runs. No model was fine-tuned on the evaluation tasks.

For model comparison, outputs were evaluated under identical task definitions and expert criteria. Response time, token consumption, classification performance and feature-level extraction quality^5^ were recorded for each model.

### Retrieval-augmented evidence grounding

For each task query *q*, LiveMat used retrieval-augmented generation rather than unconstrained prompting. The source corpus was first segmented into evidence units *C* = {*c*_*i*_} where each unit corresponded to a paragraph, table-derived record, figure-derived statement or database entry with document identifier, section location and provenance metadata. Textual and structured fields were embedded as *e*_*i*_ = *f φ* (*c*_*i*_), and the query was embedded as *e*_*q*_ = *f φ* (*q*).

Candidate evidence units were ranked by a hybrid retrieval score *R* (*q,ci*) = *λ* cos (*e*_*q*_ , *e*_*i*_) + (1 − *λ*) *BM25* (*q,c*_*i*_) , where *λ* controls the contribution of vector similarity and lexical matching. The retrieved context for a query was *C*_*q*_ = *TopK*_*i*_*R* (*q, c*_*i*_). The values of and were fixed before benchmark evaluation and were not adjusted per model or per case.

The LLM input was defined as *y* = *LLM* (*q, c*_*q*_, *p*_*task*_) where *p*_*task*_ denotes the task-specific instruction template. Outputs were accepted for downstream extraction only when they could be linked to at least one retrieved evidence unit in *c*_*q*_ or to an explicit expert-defined constraint. Statements without such support were labelled E4 and were excluded from positive graph-edge construction.

To reduce retrieval leakage, evidence retrieval, answer generation and expert adjudication were logged as separate objects. The stored trace for each output was *τ* = (*q, C*_*q*_, *y, E, a*) where E is the set of cited evidence units and a is the expert adjudication label.

### Expert annotation and participation

Expert evaluation was conducted by a panel of fifteen specialists, including five senior researchers in living materials, five microbiologists and five materials scientists. Experts assessed biological validity, material feasibility and application relevance using standardized criteria. Expert involvement was restricted to annotation, adjudication and constraint definition; experts did not edit or regenerate model outputs.

For each evaluated output, primary assessment was conducted by paired junior experts consisting of one microbiologist and one materials scientist. Cases with divergent scores, conflicting qualitative judgments or low confidence were reviewed by the senior expert panel. The consensus judgment of the senior panel was used as the reference standard. This workflow follows the evaluation structure described in the manuscript (Supplementary Fig. S2), including paired review, secondary adjudication and final consensus assessment.

### Task selection and case study design

Design tasks were defined before system execution and were selected to cover different levels of constraint complexity and data availability. Tasks were not excluded based on outcome. The acute wound-healing case study was selected because it requires simultaneous consideration of infection control, oxygen supply, tissue regeneration, material compatibility and translational feasibility.

### Expert-based classification evaluation

Classification performance was evaluated using expert-annotated L3 labels as the reference. Each sample could contain one or more labels. Model predictions were normalized by case-insensitive matching and trimming of leading and trailing spaces. Multi-label predictions were treated as unordered sets.

For a sample with expert label set *E* and predicted label set *P*, classification performance was quantified using a set-based F1 score:

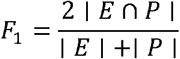

Two non-specific labels, “unrelated” and “others”, were treated as special cases. A prediction received a full score only when both the expert annotation and the model output belonged exclusively to this non-specific label set. Other samples were evaluated using the standard multi-label protocol. The final classification score was calculated as the mean F1 score across all evaluated samples.

### Expert-based feature evaluation

Feature extraction was evaluated across predefined material, microorganism and living-material dimensions. Each feature instance was assigned one of four correctness levels: Accept, Weak–Incomplete, Weak–Mixed or Reject. These categories were converted to numerical scores:

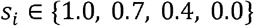

where *s*_*i*_ is the expert score for feature *i*. Feature-level accuracy was computed as the average score across all evaluated features:

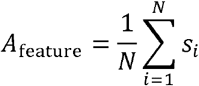

Here, *N* denotes the number of evaluated feature instances for the corresponding model and task.

Accept and weak predictions were treated as non-empty outputs for precision and recall analysis. Correct extractions were counted as true positives, unsupported or incorrect extractions as false positives, and missed expert-identified features as false negatives. Precision, recall and F1 score were calculated as:

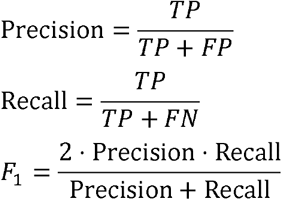

Unsupported extra-textual inference was recorded as an E4 error. The E4 error rate was defined as:

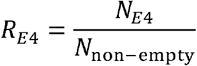

where *N*_*E*4_ is the number of E4-labeled outputs and *N* _non– empty_ is the number of non-empty model outputs.

### Evidence attribution and inference tracing

Each evaluated statement was assigned an evidence category according to its source and inference depth. Directly supported statements were labelled as direct extraction; semantically equivalent restatements were labelled as paraphrased extraction; statements requiring reasoning within the same article were labelled as in-article inference; unsupported or external claims were labelled as E4. For inferred answers, experts recorded the minimal evidence chain required to support the judgement. If no direct or indirect evidence was identified, the answer was marked as unsupported.

### Knowledge graph construction and design-space reconstruction

Extracted entities were standardized and mapped to microorganism, material, payload, delivery-system, state and attribute nodes. Relations encoded factual associations, including material–microorganism usage, encapsulation, delivery, functional role and experimental context. Additional relations represented inferred design constraints, including biosafety level, oxygen requirement, growth condition, material compatibility and processability.

For design-space reconstruction, materials and microorganisms were projected into feature-aware low-dimensional representations. Material entities were encoded using backbone chemistry, architecture, topology, loading mode and biological-function descriptors. Microbial entities were encoded using taxonomic identity, organism type, engineering status, growth features and effector-related annotations. Clustering and density analyses were used to identify dominant design regions, underrepresented regions and coverage gaps.

### Graph formalization, edge confidence and negative constraints

The reconstructed knowledge graph was treated as a typed, attributed multigraph rather than a narrative inventory. Formally, *G* = (*V, E, X*) where

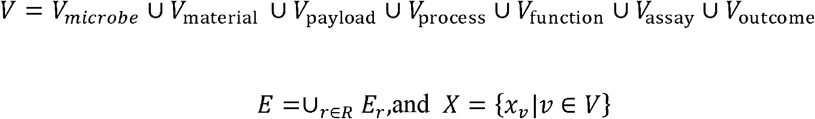

Here *R* denotes the set of relation types and *x*_*v*_ denotes the feature vector attached to node *v*.

Each node and edge were encoded with explicit provenance and evidence state:

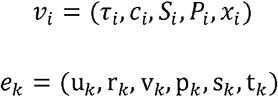

where *τ* _*i*_ is the node type, *c*_i_ is the canonical name, *S*_i_ is the synonym set, *P*_i_ is the set of source identifiers, *p*_k_ is the provenance record, *s*_k_ is the edge confidence and *t*_k_ is the evidence category.

Edge confidence was computed as a product of extraction reliability, evidence depth and cross-source consistency:

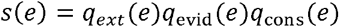

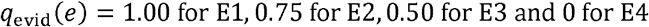

For relation type *r*, multiple pieces of evidence between the same node pair were combined by a bounded noisy-OR adjacency score, 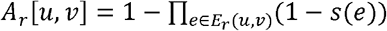. Edges with *q*_evid_ (*e*) = 0 were retained as audit records but excluded from positive ranking evidence.

Negative constraints were represented as typed inhibitory records rather than free-text notes:

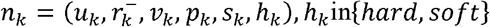

For a candidate *z*, hard and soft negative evidence were summarized as

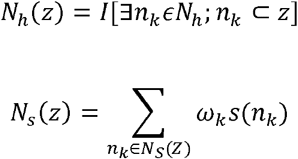

Hard negative constraints removed candidates from the feasible set, whereas soft negative constraints contributed a weighted penalty during scoring.

### Constraint-guided candidate generation from the graph

A candidate living material was parameterized as a four-component graph tuple for the wound-healing task:

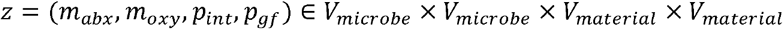

where *m*_*abx*_ denotes an antibacterial microorganism, *m*_*oxy*_ an oxygen-producing microorganism, *p*_*int*_ an interface-stabilizing matrix and *p*_*gf*_ a growth-factor-interactive matrix.

The initial candidate space *Z*_0_ was generated from graph neighborhoods satisfying the task constraint family *C* = *C*_*bin*_ ⋃ *C*_*mat*_ ⋃ *C*_*proc*_ ⋃ *C*_*app*_.

Candidate generation was therefore formulated as constraint-guided search over *Z*_0_. The feasible set was

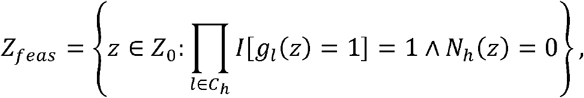

where *C*_*h*_ is the set of mandatory biological, material, process and safety constraints. Evidence-supported utility for criterion j was calculated from graph paths linking the candidate to the target function:

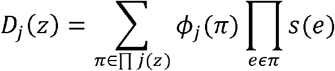

The final graph-grounded objective combined feasible utility with negative-constraint penalties:

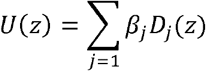

For the wound-healing case study, *β* _*j*_ was set uniformly across the seven functional dimensions unless otherwise stated, matching the equal-weight scoring used below.

### Multimodal normalization of wound-healing outcomes

Outcome measures from wound-healing studies were extracted from text, tables and figures. Metrics included wound closure, antibacterial activity, inflammation markers, collagen deposition, angiogenesis, cytocompatibility and epidermal thickness. Only studies with explicit control–treatment comparisons were included.

To compare heterogeneous metrics across studies, quantitative indicators were normalized to a common scale. For metrics where higher values indicate better performance, min–max normalization was used:

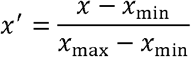

For metrics where lower values indicate better performance, inverse normalization was used:

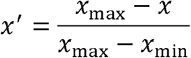

Within each functional dimension *d*, normalized indicators were averaged:

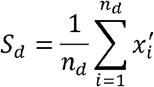

where *n*_*d*_ is the number of available indicators in dimension *d*. Missing dimensions were not imputed. The overall performance score was calculated from available dimensions:

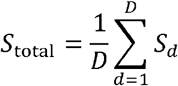

where *D* is the number of evaluated dimensions available for that system.

### Dimension-specific scoring for wound-healing evaluation

For wound closure, a stage-aware scoring model was used because healing rate depends on experimental time point. For a measured closure value *v*, a stage-specific baseline and target *T* were used:

For each sampling day, *B* and *T* were specified from the control baseline and the target closure level used for that stage, respectively; the same *B* and *T* values were applied to all systems evaluated at that time point.

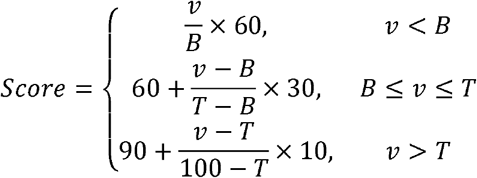

Antibacterial activity was scored according to the reported metric. For CFU-based measurements, lower bacterial load was converted to a higher score:

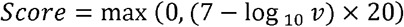

For percentage antibacterial rate, the reported percentage was used directly after truncation to the 0–100 range. For normalized CFU or survival-rate metrics where higher values indicated weaker antibacterial effect, the score was inverted as:

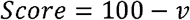

Inflammatory markers were scored according to directionality. For relative TNF-*α* or IL-6 levels, lower values indicated stronger anti-inflammatory effect:

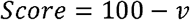

For absolute TNF-*α* concentration, the score was calculated as:

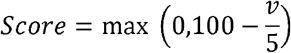

For IL-10, where higher values indicate improved anti-inflammatory response, the score was calculated as:

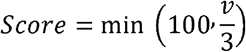

Collagen deposition was converted using:

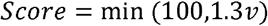

Angiogenesis was scored according to the reported indicator. For CD31-positive area, the score was:

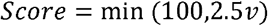

For vessel counts, the score was:

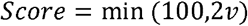

For absolute cell or vessel counts exceeding 100, group-wise relative normalization was used:

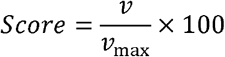

Cytocompatibility was scored directly from viability percentage:

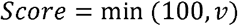

Epidermal thickness was scored against an approximate physiological optimum of 75 μm:

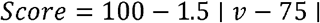

Scores below 0 were truncated to 0 and scores above 100 were truncated to 100.

### Model-proposed and agent-designed four-component systems

For the wound-healing design comparison, each large language model was prompted to propose a four-component living material system (Fig. 5F) consisting of one antibacterial microorganism, one oxygen-producing microorganism and two material components. The same task description and constraints were used across all models.

Each proposed system was evaluated across eight criteria: oxygen/carbon dioxide exchange, ROS scavenging, antimicrobial activity, immune modulation, angiogenesis support, biosafety, mechanical support and process compatibility. Each criterion was scored on a 0–10 scale by expert assessment and literature-derived evidence. The overall fitness score was calculated as (Fig. 5F):

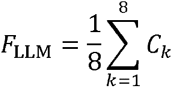

where *C*_*k*_ is the score for criterion *k*.

### Combinatorial fitness evaluation

For constraint-guided search, top-ranked oxygen-producing microalgae, antibacterial bacteria, interface-stabilizing materials and growth-factor-interactive materials were combined into a four-component design space (Fig. 5G). Three candidates were retained in each functional category, yielding 9 × 9 combinations.

Thus, 3 × 3 microbial pairs and 3 × 3 material pairs generated 9 microbial-module combinations and 9 material-module combinations, yielding an evaluated 9 × 9 design matrix.

Each combination was scored across seven dimensions: antimicrobial activity, oxygen production, moist microenvironment support, growth-factor enrichment, biocompatibility, component compatibility and absence of pharmacological contraindication. The total combinatorial fitness score was:

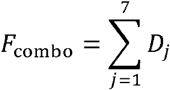

where *D*_*j*_ denotes the score of dimension *j*, each defined on a 0–10 scale. The maximum score was therefore 70.

Component compatibility was calculated from microbial–microbial compatibility, material–material compatibility and cross-domain compatibility:

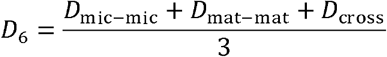

The highest-scoring combination was selected as the constrained optimum within the evaluated design space.

### Comparative ranking of literature-derived systems

Literature-derived wound-healing systems were ranked using the same normalization and scoring procedures. For each system, available dimension scores were aggregated without imputing missing values. Rankings were computed independently of the experimental results generated in this study, allowing the LiveMat-designed system to be positioned against reported living-material systems using the same evaluation scale.

### Preparation of living materials system

For preparing living microgels, calcium-crosslinked methacrylated alginate hydrogels containing different microorganisms were prepared separately. Briefly, an aqueous precursor solution containing 2 wt% methacrylated sodium alginate and 1 wt% LAP was prepared by stirring at room temperature in the dark until complete dissolution. Suspensions of *B. subtilis* and *C. vulgaris* were collected by centrifugation at OD □ □ □ = 1 and OD□ □ □ = 1, respectively. After removal of the supernatant, the cell pellets were resuspended in the methacrylated alginate precursor solution to obtain single-microorganism-loaded hydrogel precursors. The microorganism-containing precursors were slowly dropped into a 2 wt% CaCl_2_ aqueous solution to form Ca^2+^-crosslinked alginate hydrogels. After gelation, the hydrogels were collected and washed 2–3 times with sterile PBS (pH 7.2) to remove residual CaCl □. The resulting microorganism-laden hydrogels were then mechanically fragmented through a micron-scale stainless-steel mesh under aseptic conditions to obtain living hydrogel microgels (Fig. 6A).

For preparing the living materials system, two types of microorganism-laden microgels were mixed at predetermined ratios and gently homogenized to form a composite living hydrogel ink (Fig. 6A). The ink was transferred into the cartridge of a 3D printer, extruded into designed structures, and subsequently stabilized by secondary photo-crosslinking under 405 nm blue light irradiation. Hydrogels prepared using different crosslinking strategies were characterized to evaluate their viscoelastic properties, microbial distribution, and internal microstructure. Three types of samples were prepared for comparison: Ca^2+^-crosslinked hydrogels, photo-crosslinked hydrogels, and Ca^2+^/photo-dual-crosslinked hydrogels. For Ca^2+^-crosslinked samples, the methacrylated alginate/LAP precursor solution was added dropwise into a 2 wt% CaCl_2_ solution to form ionically crosslinked hydrogels, followed by washing with PBS and mechanical fragmentation through a micron-scale mesh. For photo-crosslinked samples, the precursor solution was cast into a mold and exposed to 405 nm blue light to induce covalent gelation. For dual-crosslinked samples, the Ca^2+^-crosslinked hydrogel fragments were further molded and subjected to 405 nm light irradiation to generate Ca^2+^/photo-dual-crosslinked hydrogels.

### Rheological, microscopic, and structural characterization

Rheological measurements were performed using a Thermo Scientific HAAKE MARS rheometer at 25 oC. Oscillatory time sweep tests were conducted at a frequency of 1 Hz to monitor the evolution of the storage modulus (G’) and loss modulus (G’’) of the dual-crosslinked hydrogels before, during, and after light exposure. Oscillatory amplitude sweep tests were performed over a strain range of 0.1-100% to determine the deformation-dependent viscoelastic response of hydrogels prepared by different crosslinking methods. Oscillatory frequency sweep tests were further conducted over an angular frequency range of 0.1-100 rad s-1 at a strain within the linear viscoelastic region (Fig. S16C-E) to obtain the frequency-dependent G’ and G’’ values.

The spatial distribution of microorganisms in the Ca^2^ □ /photo-dual-crosslinked hydrogels was observed using an Olympus IX73 inverted fluorescence microscope. GFP-expressing B. subtilis and red-fluorescent C. vulgaris were used to distinguish the two microbial populations within the hydrogel matrix (Fig. S17A). During image acquisition, the exposure time, gain, and other imaging parameters were kept constant for each fluorescence channel to ensure comparability among different samples.

The microstructure of the hydrogels and the distribution of embedded microorganisms were further characterized by scanning electron microscopy. Cell-laden Ca2+/photo-dual-crosslinked hydrogels were fixed overnight at 4 °C in 2.5% glutaraldehyde prepared in PBS (pH 7.2). After rinsing with PBS, the samples were dehydrated sequentially in ethanol solutions with increasing concentrations of 30%, 50%, 70%, 90%, and 100% for 15 min at each step. Ethanol was then replaced with isoamyl acetate through a 1:1 ethanol/isoamyl acetate mixture, followed by pure isoamyl acetate treatment. The samples were rapidly frozen in liquid nitrogen, lyophilized, and sputter-coated with gold before imaging. SEM observation was performed using a Hitachi TM-3000 scanning electron microscope to examine the hydrogel network morphology and microorganism encapsulation within the matrix (Fig. S17B).

### Animal

Male BALB/c mice (6-8 weeks old, 18-22 g) were obtained from Jiangsu Huachuang Xinnuo Pharmaceutical Technology Co., Ltd. (Taizhou, China). All animals were housed under specific pathogen-free (SPF) conditions with tightly controlled environmental parameters: a temperature of 22 ± 2°C, relative humidity of 50 ± 5%, and a 12 h light/dark cycle. Mice were provided with standard chow and water ad libitum and maintained in uniform cages bedded with wood shavings. All animal procedures were approved by the Institutional Animal Care and Use Committee of China Pharmaceutical University (Approval No. YSL-202604035) and were conducted in compliance with the ARRIVE guidelines. Throughout the study, efforts were made to minimize the number of animals used and to reduce any potential discomfort.

### Establish a mouse model of acute wound healing

Mice were anesthetized with isoflurane, and a full-thickness excisional wound was created at the center of the dorsal skin using a 4 mm biopsy punch, removing the epidermis and superficial dermis. A white silicone splint (inner diameter: 5 mm) was then sutured around the wound to stabilize the surrounding skin and minimize contraction. The animals were randomly assigned to four groups (n = 6 per group): (1) control group (wounds left untreated and exposed to air); (2) SA + Ca^2+^-crosslinked hydrogel group; (3) SA + CS + Ca^2+^-crosslinked + photo-crosslinked group; and (4) SA + CS + Ca^2+^-crosslinked + photo-crosslinked + *B. subtilis* + *C. vulgaris* group, which was additionally maintained under a 12 h light/12 h dark white-light regimen (8000 lux). Freshly prepared hydrogels were applied topically to the wounds in all treatment groups, while the control group received no intervention. Wound healing progression was documented by digital photography on the day of surgery day 0, day 3, and every 3 days thereafter until complete closure. Wound areas were quantified from the images using ImageJ software (Fig. 6B-D).

At the study endpoint, skin tissues were harvested from the wound area and fixed in 4% paraformaldehyde at room temperature for 24 h. Following fixation, samples were dehydrated through a graded ethanol series, cleared in xylene, and embedded in paraffin. The paraffin blocks were sectioned into 5 μm-thick slices using a microtome, and the sections were subjected to H&E staining as well as Masson’s trichrome staining according to standard protocols. Histological features were examined under an optical microscope (IX71, Olympus), and ImageJ software was used to quantitatively assess epidermal thickness (Fig. 6E,F).

### Immunofluorescent staining

CD31 immunofluorescence staining was performed using a tyramide signal amplification (TSA)-based four-color fluorescence kit according to the manufacturer’s instructions. Briefly, tissue sections were immersed in antigen retrieval buffer (either EDTA buffer, pH 9.0, or citrate buffer, pH 6.0) and subjected to heat-induced antigen retrieval using a microwave oven: sections were brought to a boil for approximately 3 min, followed by heating at medium–low power for 15–20 min, and then allowed to cool to room temperature. Endogenous peroxidase activity was quenched by incubation with 3% hydrogen peroxide for 10 min at room temperature. The sections were then blocked with 5% bovine serum albumin (BSA) for 10 min at room temperature to reduce nonspecific binding. Subsequently, the sections were incubated with anti-CD31 primary antibody (Fig. 6G; Fig. S18A) at 4°C. After washing with PBS, sections were incubated with HRP-conjugated secondary antibody for 30 min at room temperature. The fluorophore stock solution was diluted 1:100 in the provided staining diluent to prepare the working solution, followed by incubation with the sections for 10 min. Finally, sections were briefly air-dried, mounted with a DAPI-containing medium, and imaged using a fluorescence microscope (IX71, Olympus).

### Femtosecond label-free imaging (FLI)

At the experimental endpoint, skin tissue samples were harvested from the wound area and immediately fixed in 4% paraformaldehyde at room temperature for 24 h. After fixation, the samples were directly imaged without sectioning or staining using a femtosecond laser imaging (FLI) microscope system (FI-100, Femtosecond Research Center). The system delivers sub-5 fs laser pulses at a repetition rate of 12.5 MHz, with a broad spectral range of 900–1200 nm. The excitation beam is focused through a high-numerical aperture water-immersion objective (UAPON 40XW340, Olympus; NA 1.15), enabling interaction with biomolecules and structural components within the tissue for label-free, in situ imaging. Multiple nonlinear optical signals are simultaneously generated, including second harmonic generation (SHG) and third harmonic generation (THG) (Fig. 6H).

### Statistical analysis

All statistical analyses were performed using SPSS 22.0 software. Data are presented as mean ± S.D. For *in vivo* experiments, six mice were included in each group, and n represents the number of independent experiments. Statistical differences among multiple groups were evaluated by one-way analysis of variance (ANOVA), while comparisons between two groups were performed using Student’s t-test. Post hoc multiple comparisons were conducted using the least significant difference (LSD) test when variances were homogeneous and the Games-Howell test when variances were unequal. Differences were considered statistically significant at *p* < 0.05.

## Data availability

All data are available within the article and its Supplementary Information. The datasets generated and analysed during this study, including structured living materials records, model benchmarking data and source data used for computational analyses and figure generation, are available from the GitHub repository at https://github.com/YeweiXiao/Livemat1.0/tree/main/Date.

## Code availability

The code used to implement the LiveMat agent system, including the front-end interface and back-end agent workflow, is available in the GitHub repository at https://github.com/YeweiXiao/Livemat1.0/tree/main/Agent. Additional scripts used for data analysis, benchmarking and figure generation are provided within the same repository.

## Acknowledgements

This work was supported by the National Key Research and Development Program of China (2024YFA0919100) and the National Natural Science Foundation of China (T2322011, 22278214, 52003119).

## Author contributions

Y. Xiao and Z. Yu conceived the project and designed the overall study. X. Zeng designed the biological experiments, while Z. Yang designed the materials-related experiments. Y. Xiao and J. Gu developed the agent framework and associated workflows for data processing, analysis, and model integration. Y. Wang, H. Wen, Z. Huang, J. Hu, J. Liu, C. Sha, J. Xie, H. Li, X. Zhu, S. Zheng, J. Zhang, W. Zong, Z. He, Y. Xu, X. Zhou, and F. Li conducted the large language model benchmarking, data curation, feature annotation, performance evaluation, and figure generation. Y. Lu performed the biological experiments. M. Chen provided the materials and experimental resources. J. Hu carried out the materials experiments and assisted with data collection. H. Liu, Q. He, and L. Liu provided critical comments on manuscript structure, scientific presentation, and manuscript revision. Y. Xiao curated and analyzed the data and prepared the initial manuscript draft. Z. Yu, X. Zeng, Z. Yang, H. Liu, Q. He, and L. Liu revised the manuscript. All authors discussed the results, reviewed the manuscript, and approved the final version.

